# Loss of ELK1 has differential effects on age-dependent organ fibrosis and integrin expression

**DOI:** 10.1101/755694

**Authors:** Jennifer T Cairns, Anthony Habgood, Rochelle C Edwards-Pritchard, Chloe Wilkinson, Iain D Stewart, Jack Leslie, Burns C Blaxall, Katalin Susztak, Siegfried Alberti, Alfred Nordheim, Fiona Oakley, R Gisli Jenkins, Amanda L Tatler

**Affiliations:** Division of Respiratory Medicine, University of Nottingham, Nottingham University Hospitals, City Campus, Nottingham, NG5 1PB, UK and Respiratory Research Unit, NIHR Biomedical Research Centre, Nottingham University Hospitals Nottingham, UK; Newcastle Fibrosis Research Group, Institute of Cellular Medicine, Faculty of Medical Sciences, 4^th^ Floor, William Leech Building, Newcastle University, Framlington Place, Newcastle upon Tyne, NE2 4HH, UK; Department of Precision Medicine and Pharmacogenetics, The Christ Hospital Health Network, Cincinnati, Ohio, USA; Renal Electrolyte and Hypertension Division, Department of Medicine, Department of Genetics, University of Pennsylvania, Perelman School of Medicine, Philadelphia, PA, USA; Interfaculty Institute of Cell Biology, Tuebingen University, Germany and 6 Leibniz Institute on Aging (FLI), Jena, Germany

**Keywords:** Fibrosis, ELK1, ageing, TGFβ, integrin, lung, liver, gene regulation, CSE

## Abstract

ETS domain-containing protein-1 (ELK1) is a transcriptional repressor important in regulating αvβ6 integrin expression. αvβ6 integrins activate the profibrotic cytokine Transforming Growth Factor β1 (TGFβ1) and are increased in the alveolar epithelium in Idiopathic Pulmonary Fibrosis (IPF). IPF is a disease associated with ageing and therefore we hypothesised that aged animals lacking *Elk1* globally would develop spontaneous fibrosis in organs where αvβ6-mediated TGFβ activation has been implicated.

Here we identify that *Elk1*-knockout (*Elk1*^*-/0*^) mice aged to one year developed spontaneous fibrosis in the absence of injury in both the lung and the liver but not in the heart or kidneys. The lungs of *Elk1*^*-/0*^ aged mice demonstrated increased collagen deposition, in particular collagen 3α1, located in small fibrotic foci and thickened alveolar walls. Despite the liver having relatively low global levels of ELK1 expression, *Elk1*^*-/0*^ animals developed hepatosteatosis and fibrosis. The loss of *Elk1* also had differential effects on *Itgb1, Itgb5* and *Itgb6* genes expression in the four organs potentially explaining the phenotypic differences in these organs. To understand the potential causes of reduced ELK1 in human disease we exposed human cells and murine lung slices to cigarette smoke extract which lead to reduced ELK1 expression which may explain the loss of ELK1 in human disease.

These data support a fundamental role for ELK1 in protecting against the development of progressive fibrosis via transcriptional regulation of beta integrin subunit genes, and demonstrate that loss of ELK1 can be caused by cigarette smoke.

## Introduction

Tissue fibrosis is a common pathogenic pathway that results in a range of pathologies across several organs including the lung, liver, skin, kidney and heart. Fibrosis of these tissues is responsible for more than a third of deaths throughout the world [1]. There are shared, and distinct, mechanisms that may lead to fibrosis in a number of organs involving a range of cells which can provoke inflammation, epithelial injury and fibroproliferation [1,2].

A key pathway promoting fibrosis involves integrin-mediated TGFβ activation. Integrin-mediated TGFβ activation has been shown to be important in fibrosis of the lung, liver, skin, cardiovascular system and kidney [3–7] with αvβ6 integrin-mediated TGFβ activation being particularly important in the lung [4] while αvβ1 integrin-mediated TGFβ activation being considered more important for liver and kidney fibrosis [8,9].

ELK1 is an X-linked transcription factor that regulates a range of biological processes [10]. ELK1 is ubiquitously expressed in a range of cell types and in a number of organs [11]. We have previously demonstrated that ELK1 is a transcriptional repressor that prevents excessive upregulation of the β6 integrin subunit following injury [12]. Furthermore, we have shown that loss of ELK1 leads to enhanced pulmonary fibrosis following bleomycin induced lung injury in mice and that patients with Idiopathic Pulmonary Fibrosis (IPF) have reduced expression of ELK1 in their lungs [12].

Idiopathic fibrotic disease of many organs is associated with ageing. Therefore, we hypothesised that loss of ELK1 would lead to age-related tissue fibrosis of internal organs due to loss of repressed integrin expression. To test this hypothesis Elk1-knockout (*Elk1*^*-/0*^) mice were allowed to age to one year after which their internal organs were assessed for fibrosis. We demonstrated that the lungs and liver from *Elk1*^-/0^ mice developed spontaneous fibrosis whereas the heart and kidneys were normal. To understand the potential mechanism by which ELK1 may be lost in IPF, we exposed human lung epithelial cells and murine lung slices to cigarette smoke extract (CSE) and confirmed that this reduced ELK1. These studies illustrate the importance of ELK1 in protection against tissue fibrosis and highlight a mechanism through which this protective mechanism may be lost in fibrosis.

## Methods

### Cell Lines and Reagents

Immortalised human bronchial epithelial cells (iHBECs; courtesy Dr Shay, University of Texas) were used for cigarette smoke extract (CSE) experiments. iHBECs were cultured in keratinocyte-serum free medium (K-SFM; Gibco, UK) supplemented with 25µg/ml bovine pituitary extract (Gibco, UK), 0.2ng/ml epidermal growth factor (Gibco, UK), 25µg/ml G418 sulphate (VWR Life Science, UK) and 250ng/ml puromycin (MilliporeSigma, UK). ELK1 #ab32106, β-tubulin #ab6046, GAPDH #ab181602, αSMA #ab5694 and NIMP #ab2557 antibodies for western blotting and/or immunohistochemistry (IHC) were from Abcam, UK. CD3+ #MCA1477 and CD68 #OABB00472 antibodies for IHC were obtained from Bio-rad, UK and Aviva Systems Biology, UK, respectively. Reagents required for the synthesis of cDNA from RNA, were supplied from Invitrogen, UK (SuperScript™ IV Reverse Transcriptase), Qiagen, UK (Nuclease free water), Roche, UK (OligoDT) or Promega, UK (RNasin inhibitor, dNTPs). Tobacco laboratory research grade cigarettes (batch 1R6F) were from the University of Kentucky, USA.

### *In Vivo* Studies

All animal care and procedures were approved by the University of Nottingham Ethical Review Committee and were performed under Home Office Project and Personal License authority within the Animal (Scientific Procedures) Act 1986. Animals were housed in the Biomedical Services Unit, University of Nottingham. Male *Elk1*-knockout (*Elk1*^*-/0*^) [13] and wildtype (*Elk1*^*+/0*^) mice were aged for 12 or 52 weeks with free access to food (Tekland Global 18% protein rodent diet, UK) and water. Lungs, liver, kidneys and heart were either removed and snap-frozen in liquid nitrogen and then stored at -80°C for mRNA and hydroxyproline assessment, or insufflated with 10% formalin (VWR chemicals, UK) at constant gravitational pressure (20cm H_2_O) then paraffin wax embedded for histology and immunohistochemistry (IHC).

### Reverse Transcription and Quantitative Polymerase Chain Reaction (QPCR)

Murine lung, liver, kidney and heart tissue was ground into powder in liquid nitrogen and RNA extracted using TRIzol using standard methods. Alternatively, RNA from iHBECs were prepared using a NucleoSpin RNA II kit (Macherey-Nagel, UK) according to manufacturer’s instructions. For further detail see supplementary methods.

### SDS-PAGE and Western Blot

Ground murine lung, liver, kidney and heart tissue as well as Precision Cut Lung Slices (PCLS) and iHBECs were lysed in whole cell lysis buffer (Cell Signaling, UK) supplemented with protease and phosphatase inhibitors (Complete Mini protease inhibitor tablet and Phos-stop inhibitor tablets; Roche, UK) and processed for immunoblotting using standard methods see supplementary methods

### Histological Assessment

Histological sections of murine lung, liver and heart tissue, were cut to between 3-5 microns and dewaxed in xylene prior to rehydration in decreasing concentrations of ethanol.

The sections were incubated in either Mayer’s haemotoxylin, and eosin, Weigert’s haemotoxylin and Sirius red or Masson’s Trichrome prior to dehydration and mounting. Tissue staining was imaged using Nikon Eclipse 90i microscope and NIS Elements AR3.2 software (Nikon). Heart tissue sections were scanned using a Hammamatsu photonics K. K. C9600-02 and selected and captured using Hamamatsu NDP.view2 software.

### Immunohistochemistry

Tissue sections were also subjected to IHC using standard methods (see supplementary methods).

### Assessment of Tissue Fibrosis

The following scores were assigned to describe the severity of fibrosis in Sirius red stained liver sections; 0= normal, 0.5= very mild fibrosis, 1= mild fibrosis, 2= fibrosis. The following scores were assigned to assess the grade of steatosis in H&E stained liver sections; 0 = 0-5% steatosis, 1 = 5-33%, 2 = 33-66% and 3 = >66% based on the Brunt score [14]. Quantification of alveolar wall thickness in lungs of *Elk1*^*-/0*^ and *Elk1*^*+/0*^ mice was performed as follows. 10 random fields of view were observed per tissue section. Each field of view was overlaid with a graticule. The alveolar wall thickness was measured each time it intercepted a horizontal line. The mean value in µm was calculated for each mouse.

Biochemical assessment of tissue fibrosis was performed by measuring hydroxyproline in murine lung tissue as previously described [11].

### Precision Cut Lung Slices (PCLS)

6-week-old mice were euthanised their lungs were slowly inflated via a cannula to the trachea using 1.3ml 2% low melting point agarose in PBS followed by 0.2ml air. Lungs were removed and placed in ice cold DMEM supplemented with 4% L-Glutamine, 4% Penicillin/Streptomycin and 2% Amphotericin B (Gibco, UK). Lung lobes were separated and sliced in chilled HBSS buffer at a thickness of 250μm using the VT1200S Vibratome (Leica) set to 1mm amplitude, 1mm/second speed. The slices were then placed in warm supplemented DMEM media and incubated overnight at 37°C, 5% CO_2_. The slices were transferred to freezing media (FBS with 10% DMSO) and stored long-term in liquid nitrogen. PCLS were recovered by rapidly thawing frozen vials at 37°C and transferring them to fresh supplemented DMEM for a period of 24 hours at 37°C, 5% CO_2_ before use in subsequent experiments. 3 PCLS were used per condition in CSE experiments.

### Preparation of CSE and experimental design

Cigarette Smoke Extract (CSE) was generated by smoking two tobacco laboratory research grade cigarettes (batch 1R6F) into 20ml of K-SFM media (iHBEC) or DMEM (PCLS) using a vacuum pump under a constant pressure of 0.2 bar. Media was filter-sterilized and absorbance was measured at 320nm. Prepared CSE was aliquoted and stored at -20°C until required. From this 1, 2, 3, 4 and 5% CSE were prepared in K-SFM for iHBECs or DMEM (3% CSE only) for PCLS. CSE was prepared in this manner daily and applied for 7 days.

### Statistics

All data are reported as mean ± standard deviation (parametric) or median (non-parametric) of n observations. Statistical significance for parametric data was determined by either an unpaired T-test when comparing two data fields or ANOVA for comparing multiple data fields. Data that were not normally distributed were assessed by non-parametric Mann-Whitney test to determine significance. CSE experiments were repeated three times and expressed as mean data from the three independent experiments. Statistical analysis of QPCR experiments were performed using an unpaired Welch’s T-test. P values less than 0.05 were considered significant. All statistical analysis was performed using GraphPad Prism (v7.04, La Jolla, CA, USA).

## Results

### ELK1 is expressed in multiple internal organs

To determine the importance of baseline ELK1 expression in age-related fibrosis, internal organs were harvested from mice at 1 year of age followed by assessment of *Elk1* mRNA and protein. *Elk1* mRNA (**Figure 1A**) and ELK1 protein (**Figure 1B and 1C**) were expressed at the highest levels in the lung and heart tissue. In contrast, levels of *Elk1* mRNA and protein were expressed at the lowest levels in the liver. Levels of ELK1 in the kidneys were generally lower than the heart and lungs but greater than observed in the liver (**Figure 1B and 1C**).

**Figure 1:**
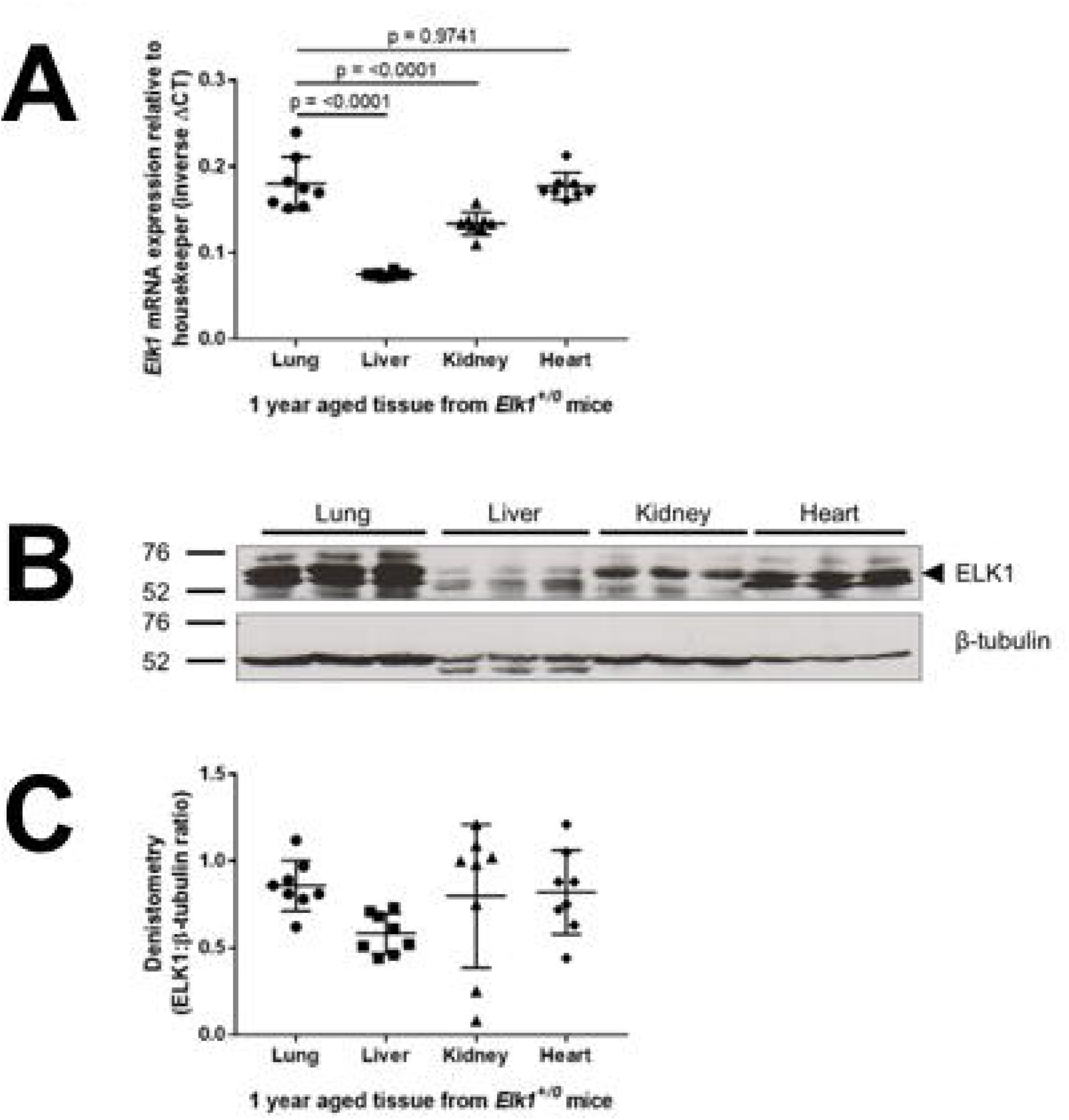
ELK1 expression in internal organs in wild-type animals. **(A)** *Elk1* mRNA expression in the lung, liver, kidney and heart tissues of 1-year aged *Elk1*^*+/0*^ mice. n=8, P values determined by one-way ANOVA. **(B)** Representative western blot showing ELK1 protein expression in the lung, liver, kidney and heart tissues of 1-year aged *Elk1*^*+/0*^ mice. **(C)** Densitometry analysis showing the ratio of ELK1:β-tubulin. n=8 mice.

### Elk1^-/0^ mice develop age related pulmonary fibrosis

The lungs from 1-year old *Elk1*^*-/0*^ mice were assessed for evidence of pulmonary fibrosis. Despite not being subjected to injury, hydroxyproline levels were mildly elevated in the lungs of *Elk1*^*-/0*^ animals compared with controls (**Figure 2A**). The elevated collagen levels in the lungs of *Elk1*^*-/0*^ mice were reflected in the lung architecture with features consistent with spontaneous lung fibrosis including small foci of alveolar destruction with increased collagen deposition and areas of lymphoid aggregates (**Figure 2B**). Analysis of these intermittent small fibrotic lesions with Sirius red staining using polarised light microscopy revealed that their composition was predominantly type I collagen (yellow/orange strand) and type III collagens (green strands) (**Figure 2C**). Furthermore, the alveolar walls of *Elk1*^*-/0*^ mice were thicker compared with wild-type control lungs (**Figure 2D and 2E**). Sirius Red staining revealed that there was increased deposition of collagen III within the alveolar interstitium (**Figure 2E**). Analysis of collagen type 1, 3 and 6 mRNA expression (**Figure 2F**) did not show any difference in *Col1a1* and *Col6a1* mRNA expression between *Elk1*^*+/0* and^ *Elk1*^*-/0*^ animals. However, *Col3a1* expression was significantly increased in *Elk1*^*-/0*^ mice consistent with the increased interstitial collagen 3 seen histologically (**Figure 2E**). Interestingly, the phenomenon of thickened alveolar walls, deposition of fibrosis and increased *Col3a1* mRNA expression was not observed in 12-week old *Elk1*^*-/0*^ mice (**Supplemental Figure 1A and 1B**), highlighting the importance of ageing in the development of this phenotype in *Elk1*^*-/0*^ mice.

**Figure 2:**
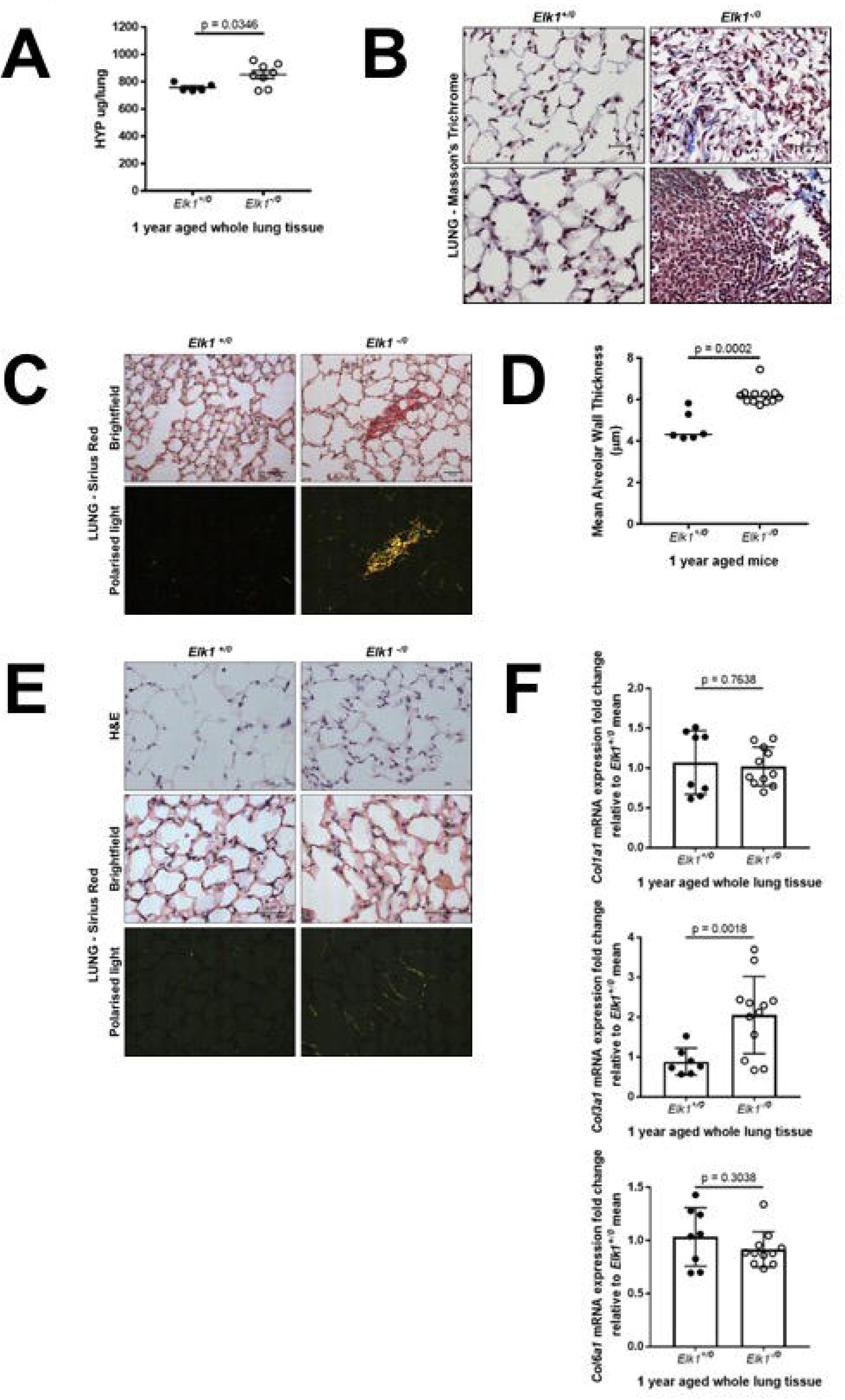
*Elk1*^*-/0*^ aged mice deposit collagen and exhibit small fibrotic lesions in the lungs. **(A)** Total lung collagen assessed by measuring hydroxyproline levels in ground lung tissue from 1-year aged *Elk1*^*+/0*^ and *Elk1*^*-/0*^ mice. P value determined by unpaired T-test. **(B)** Representative images of Masson’s Trichrome stained lung sections from 1-year aged *Elk1*^*+/0*^ and *Elk1*^*-/0*^ mice. Scale bar 25μm **(C)** Representative brightfield and polarised light images of Sirius Red stained lung sections from 1-year aged *Elk1*^*+/0*^ and *Elk1*^*-/0*^ mice. Scale bar 50μm **(D)** Quantification of alveolar wall thickness in 1-year aged *Elk1*^*+/0*^ and *Elk1*^*-/0*^ mice. P value determined by Mann-Whitney test. **(E)** Representative brightfield and polarised light images of H&E and Sirius Red stained lung sections from 1-year aged *Elk1*^*+/0*^ and *Elk1*^*-/0*^ mice. Scale bar 50μm **(F)** *Col1a1, Col3a1* and *Col6a1* mRNA expression in *Elk1*^*+/0*^ and *Elk1*^*-/0*^ in 1-year aged mice were analysed using QPCR. Data were expressed as Mrna expression fold-change relative to *Elk1*^*+/0*^ mean of 1. P value determined by T-test.

### Elk1^-/0^ mice do not develop age related cardiac or renal fibrosis

As described earlier, ELK1 is expressed at high levels in the normal heart, and to a lesser extent in the kidneys. Therefore, the heart and kidneys were assessed for phenotypic abnormalities. There was no evidence of fibrosis in the heart tissue of *Elk1*^*+/0*^ and *Elk1*^*-/0*^ animals as determined either histologically or biochemically. Indeed, there was a slight reduction of collagen in the hearts of *Elk1*^*-/0*^ animals compared with controls (**Figure 3A**), although morphological assessment of heart sections using Sirius Red showed no abnormalities in tissue structure in either wild-type or knock-out animals (**Figure 3B**). In the kidneys of *Elk1*^*+/0*^ and *Elk1*^*-/0*^ mice, there was no evidence of renal fibrosis biochemically (**Figure 3C**) or histologically (**Figure 3D**).

**Figure 3:**
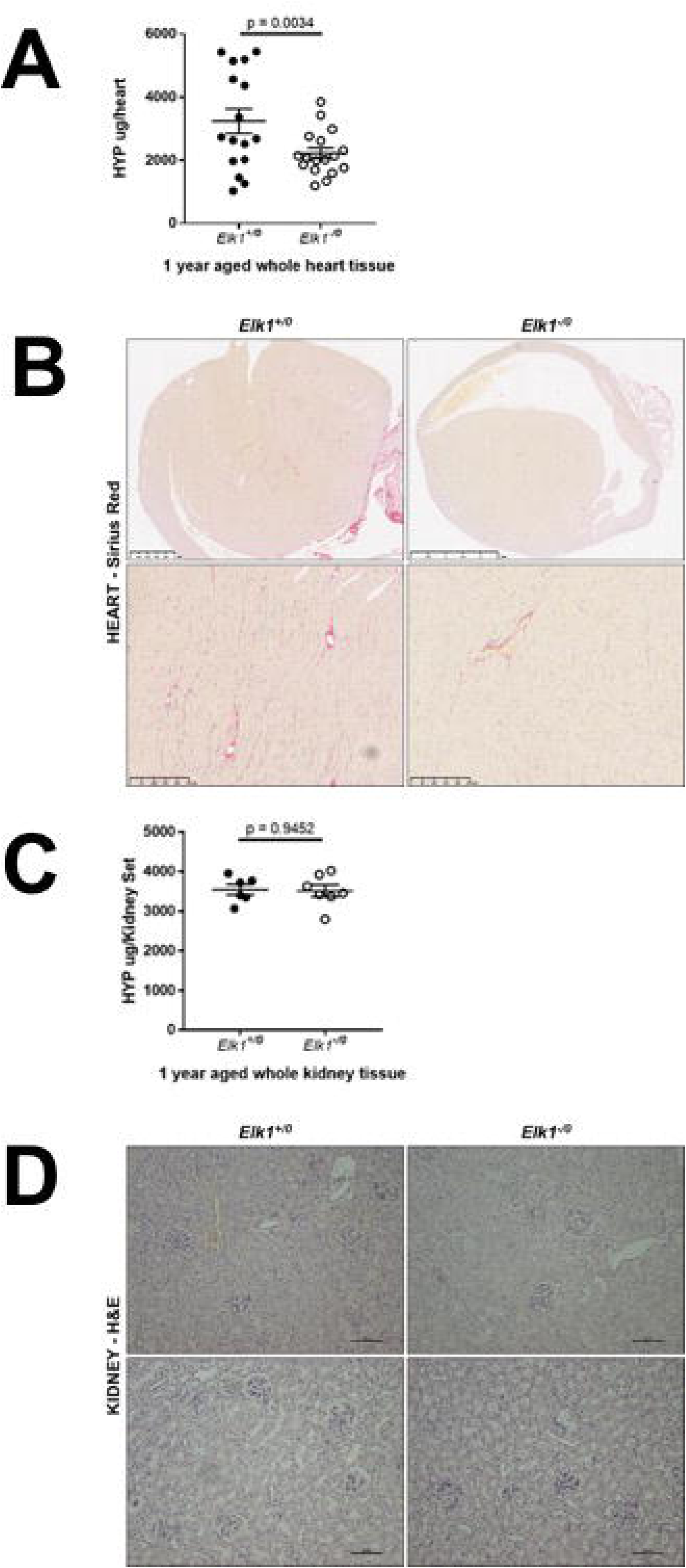
*Elk1*^*-/0*^ aged mice do not display any pathological abnormalities in the kidney or heart. **(A)** Total heart collagen assessed by measuring hydroxyproline levels in ground heart tissue from 1-year aged *Elk1*^*+/0*^ and *Elk1*^*-/0*^ mice. P value determined by unpaired T-test. **(B)** Representative images of Sirius Red stained heart sections from 1-year aged *Elk1*^*-/0*^ and *Elk1*^*+/0*^ mice. **(C)** Total kidney collagen assessed by measuring hydroxyproline levels in ground kidney tissue from 1-year aged *Elk1*^*+/0*^ and *Elk1*^*-/0*^ mice. P value determined by an unpaired T-test. **(D)** Representative images of H&E stained kidney sections from 1-year aged *Elk1*^*-/0*^ and *Elk1*^*+/0*^ mice.

### Elk1^-/0^ mice develop hepatosteatosis

Although high levels of ELK1 were not observed in whole liver lysates at either the mRNA or protein level in *Elk1*^*+/0*^ mice, there was evidence of pathology on histology, in the livers of aged *Elk1*^*-/0*^ mice. Liver sections from *Elk1*^*+/0*^ and *Elk1*^*-/0*^ aged mice, stained with Sirius Red, demonstrated increased portal fibrosis and mild perisinusoidal fibrosis in the *Elk1*^*-/0*^ animals compared with *Elk1*^*+/0*^ mice. In addition, *Elk1*^*-/0*^ mice also exhibited more micro- and macrosteatosis shown by the blue and green arrows, respectively (**Figure 4A and 4B**). However, there was no increase in hydroxyproline levels in the livers of *Elk1*^*-/0*^ mice (**Supplementary Figure 2**).

**Figure 4:**
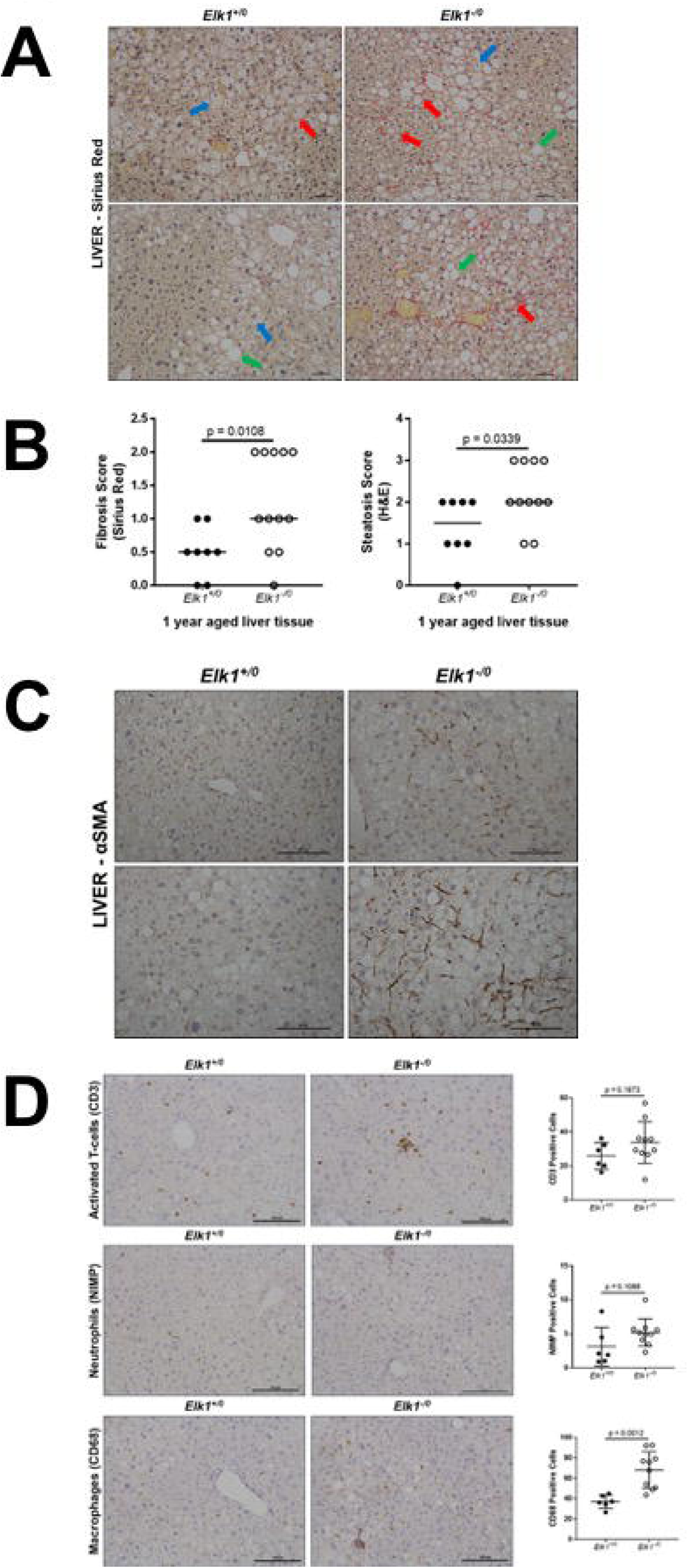
*Elk1*^*-/0*^ aged mice have mild hepatic fibrosis and steatosis in the absence of injury. **(A)** Representative images of Sirius Red stained liver sections from 1-year aged *Elk1*^*+/0*^ and *Elk1*^*-/0*^ mice showing mild fibrosis (red arrow), micro-steatosis (blue arrows) and macro-steatosis (green arrows). **(B)** H&E and Sirius Red stained liver sections from 1-year aged *Elk1*^*+/0*^ and *Elk1*^*-/0*^ mice scored blind for fibrosis and steatosis. P values determined by Mann-Whitney test. **(C)** Representative images of liver sections from 1-year aged *Elk1*^*+/0*^ and *Elk1*^*-/0*^ mice assessed for αSMA. **(D)** Representative images of liver sections from 1-year aged *Elk1*^*+/0*^ and *Elk1*^*-/0*^ mice probed for CD3 (activated T-cells), NIMP (neutrophils) and CD68 (macrophages). P value determined by unpaired T-test.

To further explore the fibrotic phenotype in the liver, immunohistochemistry (IHC) was performed using a marker of hepatic myofibroblasts. This demonstrated that alpha Smooth Muscle Actin (αSMA) was not expressed in the livers of *Elk1*^*+/0*^, however αSMA staining was increased in the livers from *Elk1*^*-/0*^ aged animals and localised to the peri-portal and peri-sinusoidal regions (**Figure 4C**).

In contrast with pulmonary fibrosis, liver fibrosis is more obviously associated with inflammation, therefore sections were assessed for the presence of inflammatory cells such as; activated T-cells (CD3), neutrophils (NIMP) and macrophages (CD68) (**Figure 4D**). Quantification of the inflammatory cell populations revealed a slight increase in the mean number of activated T-cells and neutrophils in *Elk1*^*-/0*^ mice although not statistically different. However, there was a substantial, and statistically significant, increase in the number of macrophages present in the livers from *Elk1*^*-/0*^ aged animals compared with wild-type controls (**Figure 4D**).

### Loss of Elk1 differentially affects integrin expression

To understand the phenotype of aged *Elk1* mice we assessed mRNA levels of key pro-fibrotic integrins in the lungs, liver and kidneys of *Elk1*^*-/0*^ and *Elk1*^*+/0*^ animals. In the lungs of *Elk1*^*-/0*^ mice a significant decrease in *Itgb1* mRNA was seen compared with controls animals, whereas *Itgb5* levels were unchanged between *Elk*^*+/0*^ and *Elk1*^*-/0*^ animals. There was 1.5-fold increase in *Itgb6* was observed in the lungs of *Elk1*^*-/0*^ mice compared with *Elk1*^*+/0*^ mice although this did not reach statistical significance (**Figure 5A**).

**Figure 5:**
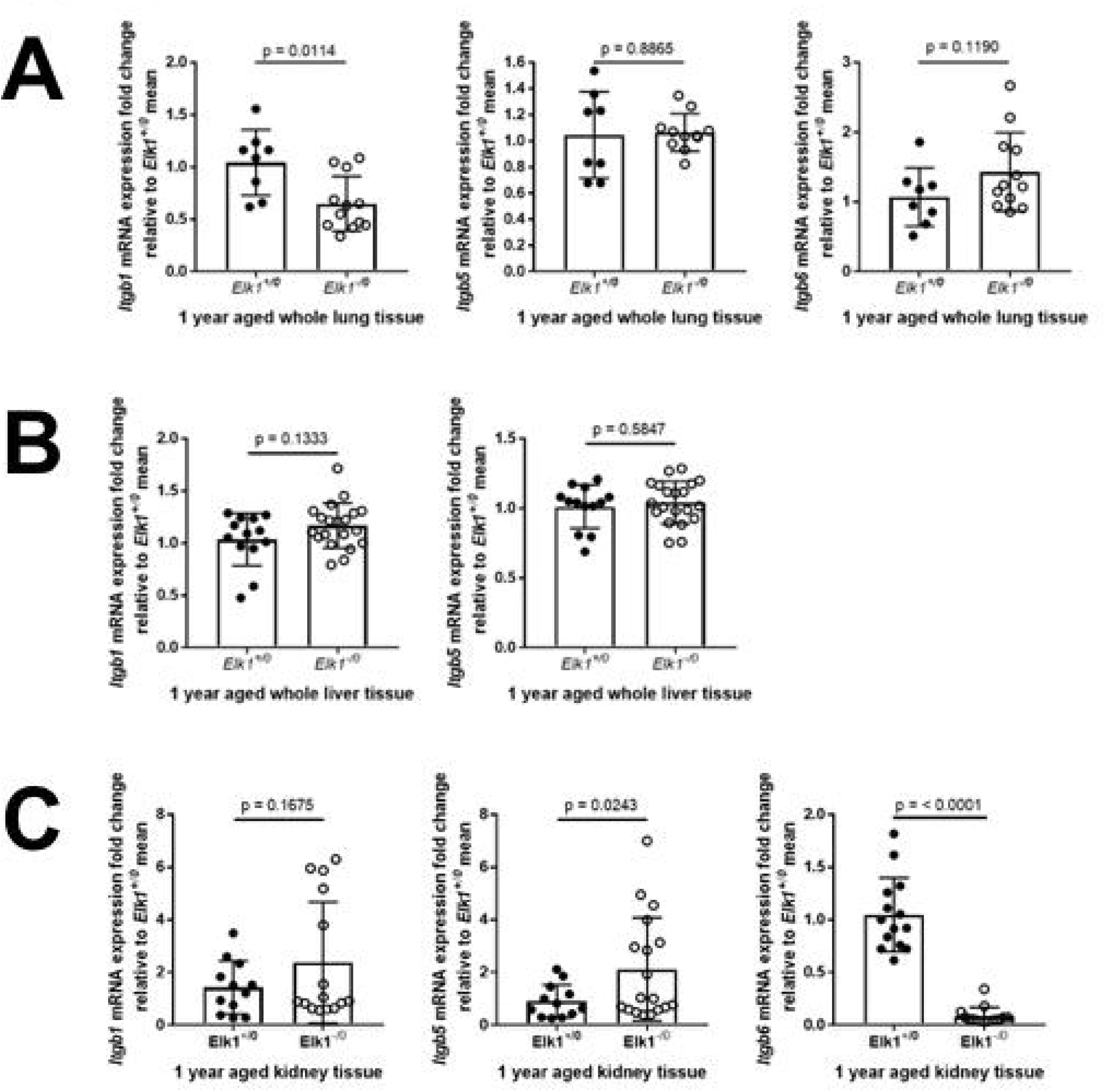
Loss of ELK1 has organ-specific effects on integrin transcription. **(A)** *Itgb1, Itgb5* and *Itgb6* mRNA expression in the lungs of 1-year aged *Elk1*^*+/0*^ and *Elk1*^*-/0*^ mice. Data were expressed as mRNA expression fold-change relative to *Elk1*^*+/0*^. P values were determined by Welch’s T-test. **(B)** *Itgb1* and *Itgb5* mRNA expression in the livers of 1-year aged *Elk1*^*+/0*^ and *Elk1*^*-/0*^ *mice we*re analysed using QPCR. Data were expressed as mRNA expression fold-change relative to *Elk1*^*+/0*^. P values were determined by Welch’s T-test. **(C)** *Itgb1, Itgb5* and *Itgb6* mRNA expression in the kidneys of 1-year aged *Elk1*^*+/0*^ and *Elk1*^*-/0*^ mice were analysed using QPCR. Data were expressed as mRNA expression fold-change relative to *Elk1*^*+/0*^. P values were determined by Welch’s T-test.

Levels of *Itgb6* were below the detection limits in the livers of both *Elk*^*+/0*^ and *Elk1*^*-/0*^ animals (data not shown) and neither whole organ expression of *Itgb1* nor *Itgb5 mRNA* expression were different between *Elk1*^*-/0*^ mice and control animals (**Figure 5B**).

The kidneys, however, showed a marked reduction in *Itgb6* mRNA levels in *Elk1*^*-/0*^ mice and conversely approximately a two-fold increase in *Itgb5* which was statistically significant and *Itgb1* levels which did not reach statistical significance (**Figure 5C**).

### Loss of ELK1 can be acquired following inhalation injury

We have previously shown that *Elk1*^*-/0*^ mice exhibit an exaggerated fibrotic response to bleomycin induced fibrosis compared with *Elk1*^*+/0*^ controls [11]. However, genetic studies have not identified an association with loss of function *Elk1* mutants and pulmonary fibrosis, and it remains unclear whether ELK1 levels can be altered following injury. Therefore, the effect of prolonged exposure to a known fibrogenic risk factor in human disease, cigarette smoke extract (CSE), on epithelial ELK1 expression was assessed.

Levels of *Elk1* mRNA in iHBECs changed in a time and concentration dependent manner following culture in CSE, with early time-points demonstrating an increase in *Elk1* particularly at higher concentrations, whereas, by day 5 exposure to CSE *Elk1* mRNA levels were reducing and by day 7 *Elk1* mRNA levels were lower than controls (**Figure 6A**).

**Figure 6:**
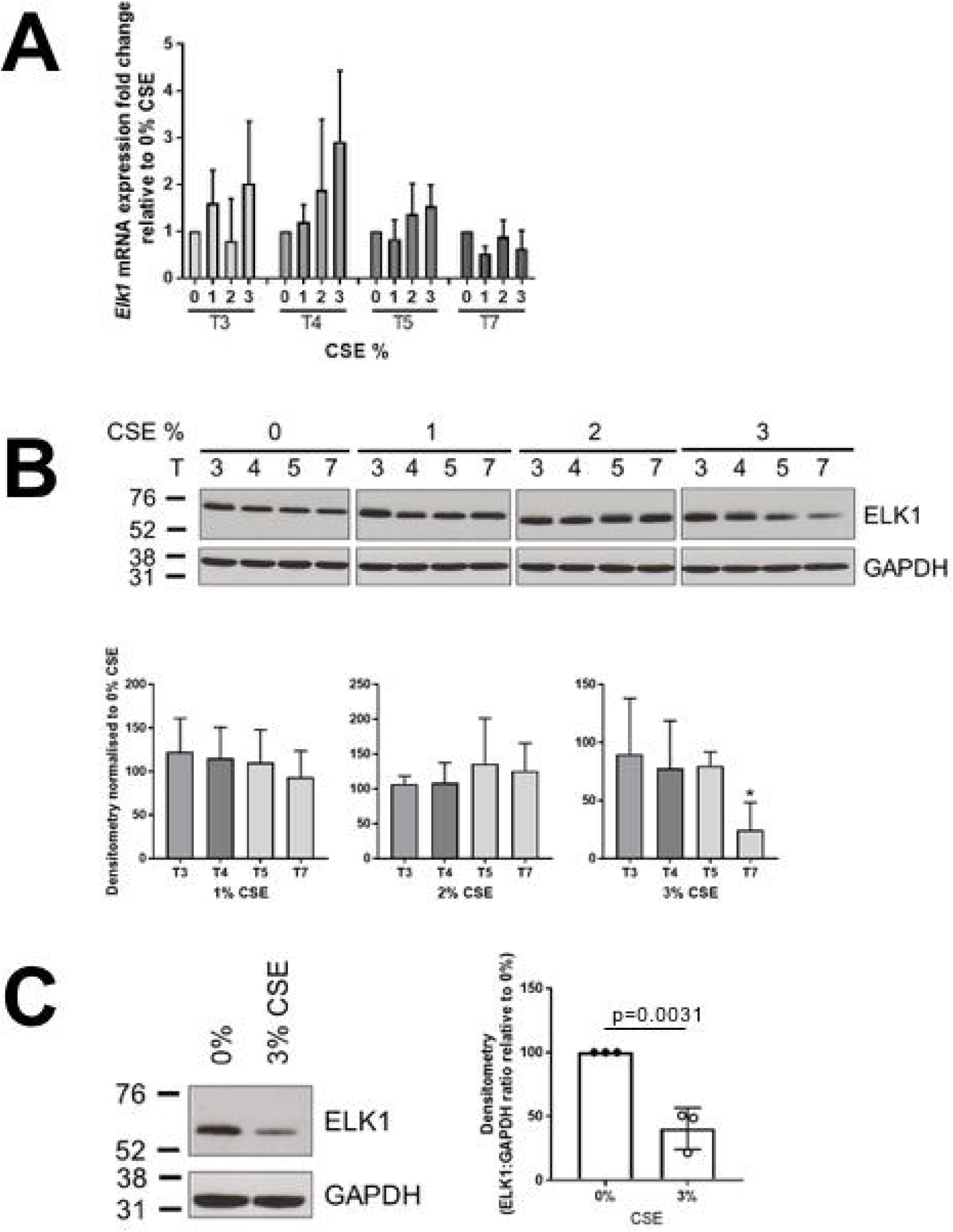
Prolonged exposure to CSE in the lungs results in the loss of ELK1. **(A)** Human *ELK1* and *GAPDH* (house-keeping gene) mRNA expression levels were analysed by QPCR in iHBECs treated with 0, 1, 2 or 3% CSE for 3, 4, 5 or 7 days (T). **(B)** Representative western blot of iHBECs treated with 0, 1, 2 or 3% CSE for 3, 4, 5 or 7 days (T). Densitometry of western blot data normalised to 0% CSE control for equivalent time point. P values determined by one-way ANOVA, 0% vs 3% CSE at T7 (n=3). **(C)** Representative western blot showing protein expression of ELK1 in PCLS treated with 3% CSE for 7 days. Densitometry of western blot data normalised to 0% CSE control. P values determined by unpaired T Test, 0% vs 3% CSE (n=3).

Western blot of lysates obtained from iHBECs exposed to 3% CSE led to reduced ELK1 after 7 days of exposure, which was confirmed by densitometric analysis (**Figure 6B**).

To confirm the results of *in vitro* cell cultures, precision cut lung slices (PCLS) obtained from uninjured lungs from wild-type mice were exposed to either 0 or 3% CSE for 7 days. Levels of ELK1 protein reduced by 50% in PCLS exposed to 3% CSE (**Figure 6C**).

## Discussion

IPF specifically, and fibrotic diseases more generally, affect primarily the elderly and have been associated with age-related processes such as telomere shortening [15–17]. We have previously identified a role for the transcriptional repressor ELK1 in the pathogenesis of IPF and have demonstrated that mice without ELK1 have an exaggerated fibrotic response to lung injury through loss of repression of αvβ6 integrin mediated TGFβ-activation [11]. We therefore hypothesised that if the ELK1 transcription repression was important for the regulation of homeostatic functions of the αvβ6 integrin then mice with global deficiency of *Elk1* would develop age-related fibrotic changes.

We assessed the phenotype of *Elk1*^*-/0*^ mice aged for 12 months without injury and found that these mice developed features consistent with pulmonary fibrosis despite the absence of injury. This is unusual in many genetic models of lung fibrosis which often require a second “hit” to promote fibrosis [18], although this may be due to the small number of studies that have looked at the effect of ageing in genetically modified mice. Some age-related studies include *Relaxin*^*-/-*^ mice that have been shown to develop features consistent with small airway fibrosis and renal fibrosis [19,20]. Furthermore, specific deletion of the telomere shelterin protein TRF1 also leads to age dependent lung fibrosis [21], as well as, mutations in genes with causal associations with IPF [22,23]. It is therefore apparent that genetic changes can promote spontaneous fibrosis across a range of organs.

It is interesting to note that the fibrotic phenotype in ELK1 deficient animals did not map onto the organs with the highest levels of ELK1 in wild-type animals. Although these data did not show a significant increase in *Itgb*6 mRNA, the direction and magnitude of change were similar to our previous studies which identified that loss of ELK1 promotes lung fibrosis through enhanced expression of the β6 integrin subunit [11]. This may explain why no phenotype relating to cardiac fibrosis was observed as epithelial injury and αvβ6 integrins are not primarily involved in fibrogenesis of the heart, where other αv integrins are involved [24].

It is interesting to note that despite the presence of considerably more fat in the livers of *Elk1*^*-/0*^ mice compared with wild-types, there was no difference in their body weight growth curves over the 1-year period (**Supplemental Figure 3**) suggesting that ELK1 has a direct hepatoprotective effect. Within the liver it is possible that ELK1 may regulate αvβ6 expression, although loss of the β6 integrin subunit did not protect against CCl4 induced hepatic fibrosis [3] and levels of *Itgb6* mRNA were undetectable in whole lung lysates in these studies. We cannot exclude that cell specific loss of ELK1 mediated repression of αvβ6 integrins may be involved in the observed phenotype, and indeed loss or inhibition of the αvβ6 integrin has been shown to protect against biliary fibrosis [5,25]. Biliary cells make up only a small proportion of the total liver cell population and it is therefore likely that any increase in cholangiocyte αvβ6 expression was not observed in the whole liver lysates used in this study. Previous studies have suggested that the αvβ1 integrin is a profibrotic integrin in both the lung and liver [3], however we consider it unlikely that ELK1 is mediating protective affects via *Itgb1* as loss of ELK lead to a reduced *Itgb1* in the lung and was unchanged in the liver, although again we cannot exclude cell specific effects.

Intriguingly, kidneys from *Elk1*^*-/0*^ mice did not show any fibrotic phenotype despite a well-established role of the αvβ6 integrin in renal fibrosis. Ischaemia reperfusion injury in rat kidneys promotes αvβ6 expression and TGFβ activation [26]. In a UUO model of renal fibrosis, mice are partially protected by loss of β6 integrin and similarly Alport mice are protected from fibrosis by administration of a β6 blocking antibody [7,27]. However, it is notable that loss of ELK1 in the kidneys had a profound effect on reducing *Itgb6* expression in the kidneys suggesting that in this organ ELK1 acts a transcriptional activator of *Itgb6*, rather than as a repressor. It is known that post-transcriptional modifications of ELK1 can affect its function [28–30]. We hypothesise that exposure to renal specific substrates may promote a distinct post-transcriptional profile, such as an excess of sumoylated ELK1, in the renal tract compared with the lung.

The role of the αvβ1 integrin in renal fibrosis is highly cell specific with deletion of β1 in collecting duct principle cells promoting fibrosis [31] but, loss of fibroblast αvβ1 integrins ameliorates it [3,9]. The data presented here show a mild and insignificant increase in *Itgb1* mRNA expression which may explain the lack of fibrosis in this aged model. Unlike the lung and the liver, *Itgb5* is increased in the kidneys of *Elk1*^*-/0*^ mice at the message level compared to controls and increased myofibroblast expansion has been associated with increased αvβ5 levels in the kidney [32]. It is clear from our studies that there are cell and organ specific effects of ELK1 on the regulation of the different integrin subunits which are likely to determine the fibrotic phenotype associated with ageing.

This study provides detailed phenotyping of uninjured age-related fibrosis in *Elk1*^*-/0*^ mice, including the identification of ELK1 as an age dependent mediator of lung and liver fibrosis in male C57BL/6 mice. As the *Elk1* gene is located on the X chromosome, there are particular benefits to using only male mice in these studies including eliminating the effects of female hormones on the fibrotic response [33]. Whilst ageing the mice beyond one year may have further exaggerated the age-related fibrotic effects and given more dramatic phenotypes it is reassuring that a phenotype was observed after only 12 months. A further strength is the application of a known fibrosis risk factor on both a cell line and physiologically viable PCLS both demonstrating a link between smoking and the loss of ELK1.

In conclusion, our data support a role for the loss of ELK1 in promoting age-related fibrosis in both the lung and liver. These data also highlight the organ-specific effects of ELK1, including a differential effect on *Itgb6* in the lung and kidney, where ELK1 appears to act as a transcriptional activator of *Itgb6* gene expression. Finally, exposure to known fibrogenic risk factors can reduce expression of ELK1 in the lung epithelium, which may in part explain how cigarette smoke increases the risk of IPF. These findings highlight the importance of normal functioning ELK1 in protecting two major organs from age-related fibrosis.

## Acknowledgements

We would like to acknowledge the Nuffield Foundation and the two placement students, Miss Zainab Mubashra and Miss Gurleen Virk, whose input has supported this project. Also, we would like to thank Miss Rachel Walker, her efforts as a BSc student have aided this work. Finally, we thank the Nottingham Health Science Biobank (NHSB) for help with tissue processing and the School of Life Sciences IMaging (SLIM) for their microscopy expertise. AN was supported by the DFG (SFB/TR 209, Liver cancer). RGJ is supported by an NIHR Research Professor award (NIHR-RP-2017-08-ST2-014). ALT received support for this work from MRF-091-0004-RG-TATLE and MRFAUK-2015-312.

## Author contributions

RGJ and ALT conceived, designed, and supervised the study. JTC, AH, FO, CW and JL performed experiments. AN and SA were involved with studies using *Elk1*^*-/0*^ mice. JTC and IDS performed data analyses. JTC, BB and KS provided histopathological assessment. JTC, AH and RCEP facilitated collection and processing of mouse tissues. JTC, RGJ and ALT drafted the manuscript. All authors provided comments on the manuscript and had final approval of the submitted version.

## Supplementary material

### Methods

#### Reverse Transcription and Quantitative Polymerase Chain Reaction (QPCR)

RNA was reverse transcribed into cDNA using SuperScript™ IV Reverse Transcriptase. cDNA was subjected to QPCR analysis using Kapa Taq to assess gene expression of both murine and human genes. This was measured using an MxPro3000 or MxPro3005 instrument (Stratagene, CA, USA). Murine β2-microglobulin (B2m) and human GAPDH were used as housekeeping genes. All QPCR reactions were performed at an annealing temperature of 60°C with the exception of murine Col6a1 at 59oC. Amplification of a single DNA product was confirmed by melting curve analysis. Data were presented as expression fold-change relative to Elk1+/0 mean of 1 using the ΔΔCt equation. Gene primer sequences used are as follows: murine β2-microglobulin (*B2m*) forward 5’-CACCCGCCTCACATTGAAAT-3’ and reverse 5’-CTCGATCCCAGTAGACGGTC-3’, murine *Elk1* forward 5’-TCCCCACACATACCTTGACC-3’ and reverse 5’-ACTGATGGAAGGGATGTGCA-3’, murine *Col1a1* forward 5’-AGCTTTGTGGACCTCCGGCT -3’ and reverse 5’-ACACAGCCGTGCCATTGTGG-3’, murine *Col3a1* forward 5’-AGGTACCGATTTGAACAGGCT-3’ and reverse 5’-TTTGCAGCCTGGGCTCATTT-3’, murine *Col6a1* forward 5’-GATGAGGGTGAAGTGGGAGA -3’ and reverse 5’-CAGCACGAAGAGGATGTCAA-3’, murine *Itgb1* forward 5’-TGTGGGGAGTGGGTAGT -3’ and reverse 5’-GACACTTTGCCAACCCA-3’, murine *Itgb5* forward 5’-CCAATGCCATGACCATC -3’ and reverse 5’-CATTTCATAACGGGCCC-3’, murine *Itgb6* forward 5’-TCTGAGGATGGAGTGCTGTG -3’ and reverse 5’-GGCACCAATGGCTTTACACT - 3’, human *GAPDH* forward 5’-ACAGTTGCCATGTAGACC-3’ and reverse 5’-TTTTTGGTTGAGCACAGG-3’, human *ELK1* forward 5’-CCACCTTCACCATCCAGTCT-3’ and reverse 5’-TCTTCCGATTTCAGGTTTGG-3’ and human *ITGB6* forward 5’-TGGGGATTGAACTGCTTTGC-3’ and reverse 5’-AGGTTTGCTGGGGTATCACA-3’.

#### SDS-PAGE and Western Blot

Briefly, 100µg of tissue lysate or 50µg of cell lysate were loaded onto a 10% SDS-PAGE gel and subject to electrophoresis for 2 hours 15 minutes at 150V. The gel was transferred onto a PVDF membrane for 4 hours on ice at 110V. The membrane was blocked with 5% non-fat milk in 0.1% Tween 20 in TBS, then probed with the following antibodies for 16 hours at 4°C: rabbit anti-ELK1, rabbit anti-β-tubulin, rabbit anti-GAPDH. The membrane was incubated with western blotting detection reagent Clarity™ (Biorad, UK) for ELK1 or ECL (Amersham, UK) for GAPDH and β-tubulin and exposed to Hyperfilm-ECL (Amersham, UK) to visualise bands. Western blot films were electronically scanned and band density calculated using ImageJ.

#### Immunohistochemistry

Briefly sections underwent antigen retrieval prior to incubation with primary antibodies: anti-ELK1, anti-αSMA, anti-NIMP, anti-CD3+or anti-CD68 overnight at 4°C. Staining was visualised with DAB. All staining was captured using a Nikon Eclipse 90i microscope and NIS Elements AR3.2 software (Nikon). Cell counts were performed on liver sections that were processed for IHC and probed for CD3+, NIMP and CD68. 10 (CD3+ and NIMP) or 15 (CD68) fields of view were selected at random and positively stained cells were counted. An average was calculated for each sample and data combined.

---

**Figure S1:** Collagen levels in the lungs of 12-week aged *Elk1*^-/0^ mice are not altered

(A) *Col1a1, Col3a1* and *Col6a1* mRNA expression in 12-week aged *Elk1*^+/0^ and Elk1-/0 mice were analysed using QPCR. Data were expressed as mRNA expression fold-change relative to *Elk1*^+/0^. P values were determined by Welch’s T-test.

(B) Representative brightfield and polarised light images of H&E and Sirius red stained sections of lung tissue from 12-week *Elk1*^-/0^ and *Elk1*^+/0^ mice.

**Figure S2:** Hydroxyproline levels in the livers of 1-year aged *Elk1*^-/0^ mice are not altered

Total liver collagen from 1-year aged *Elk1*^+/0^ and *Elk1*^-/0^ mice. P value was determined by an unpaired T-test.

**Figure S3:** Growth rates are comparable in *Elk1*^+/0^ and *Elk1*^-/0^ mice

Weight in grams of C57BL/6 *Elk1*^+/0^ and *Elk1*^-/0^ mice from 5 to 52 weeks.

