## Supplementary figures and images for "Loss of ELK1 has differential effects on age-dependent organ fibrosis and integrin expression"

### Supplemental fig 3

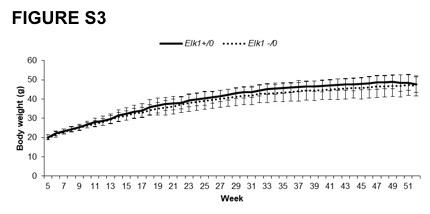

### Supplemental figure 1

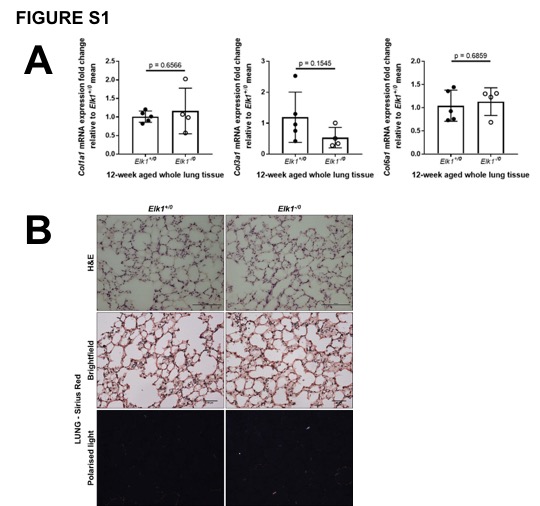

### Supplemental figure 2

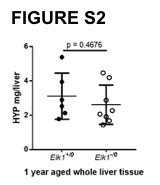
